# A λ-dynamics investigation of insulin *Wakayama* and other A3 variant binding affinities to the insulin receptor

**DOI:** 10.1101/2024.03.15.585233

**Authors:** Monica P. Barron, Jonah Z. Vilseck

**Author notes:** CORRESPONDING AUTHOR: Jonah Z. Vilseck (Address: 410 West 10^th^ Street, Suite 5017, Indianapolis, IN, 46202).

## Abstract

Insulin *Wakayama* is a clinical insulin variant where a conserved valine at the third residue on insulin’s A chain (Val^A3^) is replaced with a leucine (Leu^A3^), impairing insulin receptor (IR) binding by 140-500 fold. This severe impact on binding from such a subtle modification has posed an intriguing problem for decades. Although experimental investigations of natural and unnatural A3 mutations have highlighted the sensitivity of insulin-IR binding to minor changes at this site, an atomistic explanation of these binding trends has remained elusive. We investigate this problem computationally using λ-dynamics free energy calculations to model structural changes in response to perturbations of the Val^A3^ side chain and to calculate associated relative changes in binding free energy (ΔΔ*G*_bind_). The *Wakayama* Leu^A3^ mutation and seven other A3 substitutions were studied in this work. The calculated ΔΔ*G*_bind_ results showed high agreement compared to experimental binding potencies with a Pearson correlation of 0.88 and a mean unsigned error of 0.68 kcal/mol. Extensive structural analyses of λ-dynamics trajectories revealed that critical interactions were disrupted between insulin and the insulin receptor as a result of the A3 mutations. This investigation also quantifies the effect that adding an A3 C_δ_ atom or losing an A3 C_γ_ atom has on insulin’s binding affinity to the IR. Thus, λ-dynamics was able to successfully model the effects of subtle modifications to insulin’s A3 side chain on its protein-protein interactions with the IR and shed new light on a decades-old mystery: the exquisite sensitivity of hormone-receptor binding to a subtle modification of an invariant insulin residue.

**SIGNIFICANCE STATEMENT:** This work addresses a decades-old question of how subtle modifications to insulin’s A3 side chain affects its binding affinity to the insulin receptor. λ-Dynamics computed free energies of binding match experimental activity trends with high accuracy. Atomistic insights into hormone-receptor protein-protein interactions were obtained through a detailed investigation of λ-dynamic trajectories. This work quantifies the effects of adding and removing atoms to insulin’s conserved A3 residue and identifies clear conformational preferences for insulin A3 residues when bound to the insulin receptor.

## INTRODUCTION

As an essential component of metabolic regulation throughout the body, insulin is, perhaps, best known in the context of diabetes and the deleterious impact on health that results from the body’s failure to either produce insulin (causing type 1 diabetes) or appropriately respond to its presence (leading to type 2 diabetes). Impressively complex for its small size, this 51-residue protein is made up of two distinct chains with inter- and intrachain disulfide bonds, three α-helices, and a β-strand (1). Secreted as a hormone in response to glucose, insulin binds to the cell surface-bound insulin receptor (IR) (2), a tyrosine kinase receptor made up of two disulfide-linked homodimers that each contain an insulin-binding α subunit and the kinase-containing β subunit (3–7). Unlike most tyrosine kinase receptors, however, the IR dimers are covalently linked and remain associated even in an inactive state (3, 8–11). Insulin binding to the IR induces domain rearrangements and brings the two IR intracellular kinase domains together for auto-phosphorylation and activation of downstream signaling through the mitogenic MAPK pathway or the metabolic PI3K/AKT pathway (12–19). As a result, IR’s metabolic signaling in the liver, adipose tissue, and skeletal muscle increases glucose uptake, shuts down gluconeogenesis and lipolysis pathways, and increases glycogen, fatty acid, and amino acid synthesis. Extensive reviews describe this in greater detail (19–25).

Recent cryo-EM structures of insulin bound to the IR reveal that the IR has two sets of identical binding sites, referred to as sites 1 and 2, with higher and lower affinity for insulin respectively (Fig. 1) (26–31). At site 2, insulin primarily interacts with the FnIII-1 domain of one dimer and has minimal interactions with the L1 domain of the other dimer. Consequently, insulin binding at site 2 is not essential for IR activation (32). However, when insulin binds at site 1, it activates IR domain rearrangements and initiates a signaling cascade by interacting with the αCT peptide and a loop of the FnIII-1 domain from one IR dimer and the L1 domain of the other. This is accomplished by threading insulin’s B chain C-terminus (B^Cter^) between the one dimer’s L1 domain and the other dimer’s αCT peptide, allowing for about half of insulin’s 51 residues to directly contact the IR (33), as observed in several structures (11, 15, 26, 27, 31, 34–35). Many of these contacting residues are highly conserved and important for insulin-IR binding, as identified in a comparison study of insulin sequences across 60 different species (covering mammals, reptiles, birds, amphibians, and fish) (39). That work revealed that 16 insulin residues were completely conserved across all species, and an additional six residues were conserved in at least 55 out of the 60 species (Fig. S1) (39). Mutation of a conserved IR-interacting insulin residue would therefore be expected to negatively impact insulin’s binding interactions with the IR. For example, the first three insulin variants clinically identified in patients, named *Chicago* (Phe^B25^Leu) (40, 41), *Los Angeles* (Phe^B24^Ser) (42–44), and *Wakayama* (Val^A3^Leu) (45–49) were found to alter insulin’s binding affinity for the insulin receptor without significantly altering insulin’s independent folding or processing (40, 44, 45, 50, 51). Notably, these three residues are highly conserved across all species (Fig. S1) (39) and were later shown to directly contact the receptor (52, 53). Collectively, insulin *Wakayama*’s Leu^A3^ mutation displayed the lowest binding affinity for the insulin receptor, with 140-to 500-fold worse binding affinity relative to native Val^A3^ insulin (46, 54, 55).

**Figure 1.**
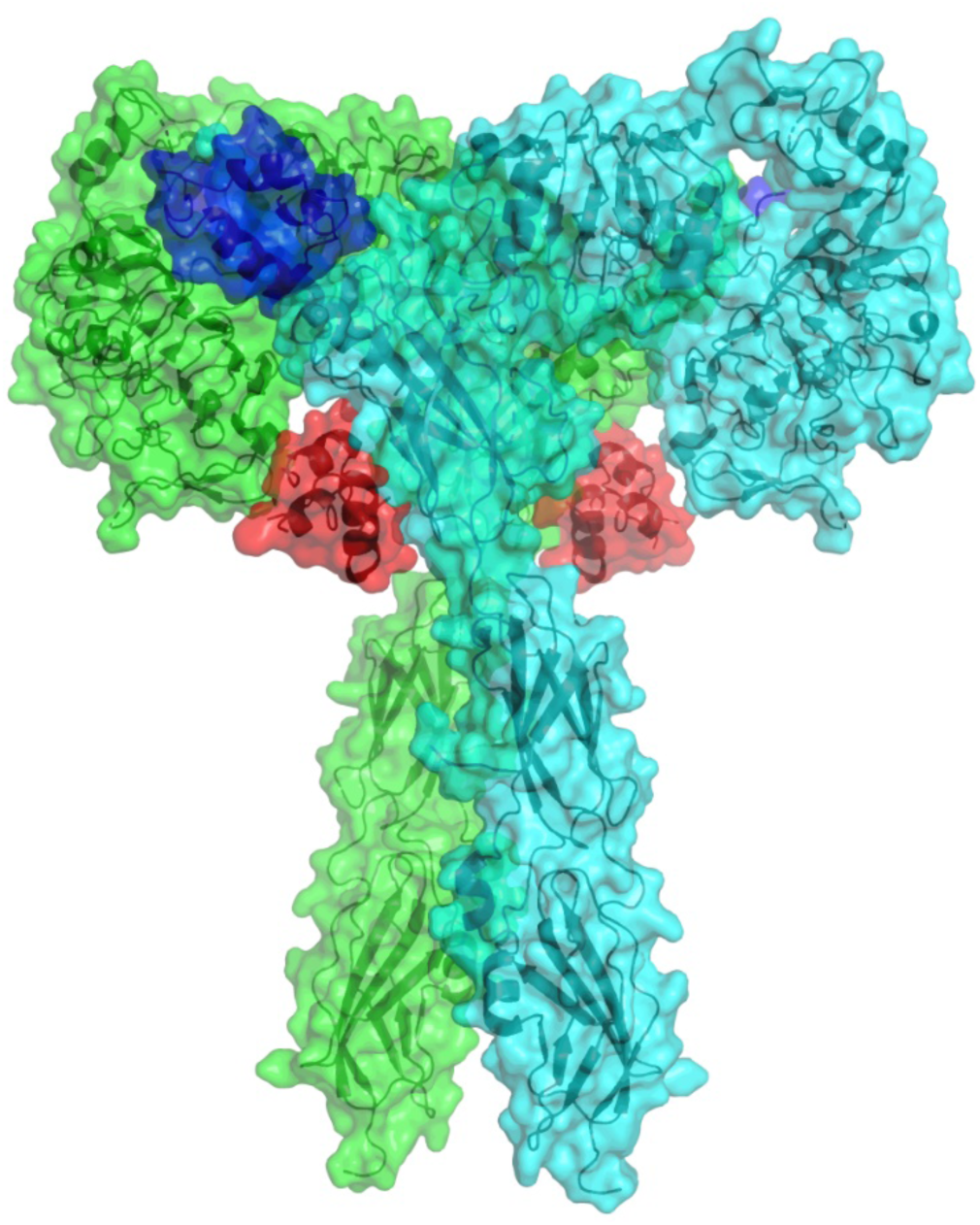
Fully bound insulin - insulin receptor ectodomain (PDBID: 6PXV (26)). The two IR homodimers are shown in green and cyan. Insulins are bound at IR sites 1 and 2, shown in dark blue and red, respectively.

To understand how the subtle addition of a single carbon unit (CH_n_) at the A3 site could produce such a drastic worsening of receptor affinity, in 1992, Nakagawa and Tager investigated IR binding affinities of many A3-modified insulin variants (56). They replaced Val^A3^ with amino acids leucine (Leu), isoleucine (Ile), alanine (Ala), and threonine (Thr) and also tested non-natural, synthetic aliphatic residues *allo*-isoleucine (Ail), *tert*-leucine (Tle), α-aminobutyric acid (Abu), and norvaline (Nva) (56). In this study, Val’s β-branching was specifically noted as important for receptor binding. Unbranched A3 residues had binding affinities of approximately two orders of magnitude worse than Val^A3^. In contrast, other β-branched A3 side chains generally produced only a single order of magnitude worse binding affinity than Val^A3^. Leu^A3^, with its γ-branched sidechain, had the most detrimental impact on insulin’s receptor binding affinity with more than a 500-fold drop in binding affinity, substantially more than its β-branched stereoisomers Ile^A3^, Ail^A3^, and Tle^A3^. Nakagawa and Tager postulated that A3 branching at the C_γ_ atom made the Leu^A3^ residue a bad fit for the receptor and that this produced insulin *Wakayama*’s greatly decreased binding affinity (56).

Due to previous difficulties in acquiring structures of the insulin-IR complex, computational studies modeling insulin-IR binding are still new, and previous structural investigations into insulin *Wakayama* have focused on insulin *Wakayama*’s hexameric and monomeric states, rather than the insulin-IR complex. The first of these studies, published in 2005, solved the crystal structure of the insulin *Wakayama* hexamer, revealing that the hexamer structure is consistent between Val^A3^ and Leu^A3^ insulin (57). Another study, published in 2017, performed molecular dynamics (MD) simulations of the native insulin monomer in solution as well as six insulin variant monomers, including insulin *Wakayama*. The data from this study revealed that, compared to Val^A3^ insulin, Leu^A3^ insulin produced no significant change to the hydrophobic interactions within insulin’s core, displayed similar conformational changes and C_α_ root-mean square deviations (RMSD) to native insulin, and showed only a modestly decreased frequency (23 % vs. 27 %) of the B^Cter^ moving into the detached “active” conformation of insulin (58). Thus, these studies suggest that Leu^A3^ does not significantly alter insulin’s conformation, dynamics, and energetics in its monomeric unbound state, further supporting the theory that the loss of binding from insulin *Wakayama* comes from a direct clash with the receptor. Despite significant advancements over the last 30 years in understanding insulin’s interactions with the IR, Nakagawa and Tager’s hypothesis that insulin *Wakayama*’s weaker IR binding is due to Leu^A3^’s γ-branching remains unproven. However, with recent cryo-EM structures of the insulin-IR complex now available, rigorous computational investigations of this problem can be performed to probe and address this question at an atomistic level.

Computational methods, such as molecular dynamics and alchemical free energy calculations enable the dynamic modeling of native and variant protein complexes. With λ-dynamics (λD), an MD-based alchemical free energy method, chemical modifications can be introduced into molecular complexes and changes in relative binding free energies (ΔΔ*G*_bind_) that result can be quantitatively computed (59–63). The thermodynamic cycle in Fig. 2 shows how free energy calculations can be used to determine ΔΔ*G*_bind_ of a mutated protein-protein complex. Unlike traditional alchemical free energy methods, λD uses a dynamic coupling parameter (λ) to investigate protein side chain mutations, and two or more chemical end states can be sampled simultaneously within a single MD simulation. This can significantly reduce overall computational costs of running these rigorous calculations. λD has been previously shown to be effective for investigating chemical modifications to either small molecule ligands, to compute relative changes in binding affinities, or protein side chain mutations, to compute protein folding stabilities (64–69). Recently, our lab has also investigated the effects of missense mutations on protein-protein binding with λD, a first for this technique (62). Specifically, 3 missense mutations of Pup2 (an α subunit) were observed to destabilize binding within the yeast 20S proteasome. Experimental stability trends were successfully reproduced by multisite λD modeling, and structural analyses revealed surprising epistasis between concurrent mutations (70), confirming that λD could successfully model protein side chain mutations and capture changes in binding affinities within protein-protein complexes. Similarly, another study investigated the effects of side chain mutations on peptide-substrate binding affinities to a protein lysine methyltransferase called PR domain zinc finger protein 9 (PRDM9). That work revealed the λD is also well suited for characterizing peptide-protein binding interactions which high accuracy compared to experiment (71).

**Figure 2.**
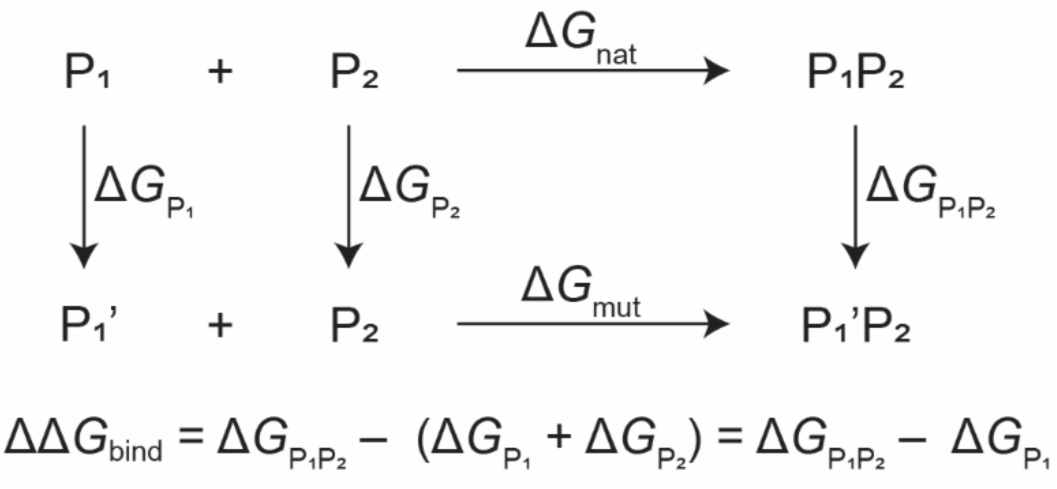
A thermodynamic cycle used to compute relative binding free energies (ΔΔ*G*_bind_) of a protein-protein complex, when one protein (P1) is mutated into a new state (P1’).

Building upon our recent successful analyses of protein-protein interactions (PPIs), in this study, we utilize λD simulations to model and explain the loss of IR binding associated with insulin *Wakayama*. Leu^A3^ and many of the A3 variants investigated by Nakagawa and Tager are modeled to explain how changes at the A3 position affect insulin’s binding affinity to the IR. A deeper understanding of Leu^A3^ insulin’s loss of IR binding is achieved via comparisons to various A3 side chains that feature changes in alkyl chain length and stereoisomerization accompanied by corresponding changes in binding magnitude. Following the successful reproduction of A3 binding thermodynamics, the λ-dynamics trajectories are extensively investigated. Preferred orientations of each insulin A3 variant are identified by side chain dihedral clustering. Mutation impacts on local insulin-IR interactions are quantified via insulin-IR distance measurements and decomposed into a per-carbon-unit of binding free energy. Collectively, this work confirms that loss of binding for insulin *Wakayama* is rooted in its binding thermodynamics with the IR, and that Leu^A3^’s second C_δ_ atom sterically clashes with the IR’s αCT peptide, which breaks multiple insulin-IR interactions and significantly reduces insulin’s IR binding affinity. Thus, λD confirms Nakagawa and Tager’s longstanding hypothesis about Leu^A3^ binding and provides new atomistic insights into insulin’s recognition by the IR at insulin’s A3 position.

## RESULTS

Insulin-IR interactions were probed as a function of insulin A3 side chain identity with λD. The scalability of λD enables multiple A3 mutations to be analyzed simultaneously, and, in practice, modeling 5-6 physical end states per λD simulation has provided a “sweet spot” for balancing computational efficiency and accuracy. Therefore, three sets of λD calculations were performed in this work to investigate the Leu^A3^ *Wakayama* mutation and seven additional A3 mutations. The first λD mutation set (Set 1) contains only the mutation of native Val^A3^ to insulin *Wakayama* Leu^A3^. The second set (Set 2) models Leu^A3^ and its stereoisomers Ile^A3^, Ail^A3^, and Tle^A3^. Finally, five variants were analyzed in the third set (Set 3): Leu^A3^, Thr^A3^, and the unbranched amino acids Ala^A3^, Abu^A3^, and Nva^A3^. The latter two represent unbranched versions of Val and Leu, respectively. By including Leu^A3^ in each calculation set, we provide a positive control to ensure all calculations are adequately converged and comparable to a consistent Val^A3^ reference state.

Prior to modeling Sets 2 and 3 with λD, the Leu^A3^ *Wakayama* mutation in Set 1 was evaluated at IR sites 1 and 2 to determine if one or both binding sites were involved in Leu^A3^’s loss of IR binding. When insulin binds the IR at site 1, one of its Val^A3^ C_γ_ atoms is partially solvent exposed while most of the residue sits in a hydrophobic pocket between Asp707, His710, and Asn711 on the IR (26, 27, 33). In contrast, at site 2 on the IR, insulin’s Val^A3^ does not contact the IR and appears completely solvent exposed (26, 27). Based on structure alone, we hypothesized that A3 mutations would only affect binding affinity at IR binding site 1. This was evaluated by performing 15 ns multisite λD simulations to interconvert between Val^A3^ and Leu^A3^ residues on four insulins simultaneously bound to a single IR (see Methods for computational details). This initial investigation is referred to as “Full”, signifying that a fully saturated IR complex was modeled. As shown in Table 1, a modest 22-fold drop in binding affinity was calculated for Leu^A3^ at binding site 1 (ΔΔ*G*_bind_ 1.8 ± 1.0 kcal/mol, “Full site 1”), but no statistically significant change in binding affinity was observed for Leu^A3^ binding at site 2 (ΔΔ*G*_bind_ -0.4 ± 0.9 kcal/mol, “Full site 2”). This confirmed that the most detrimental A3 insulin modification only altered binding to IR site 1. Therefore, all further λD simulations focused on characterizing insulin binding to site 1 only.

**Table 1.**
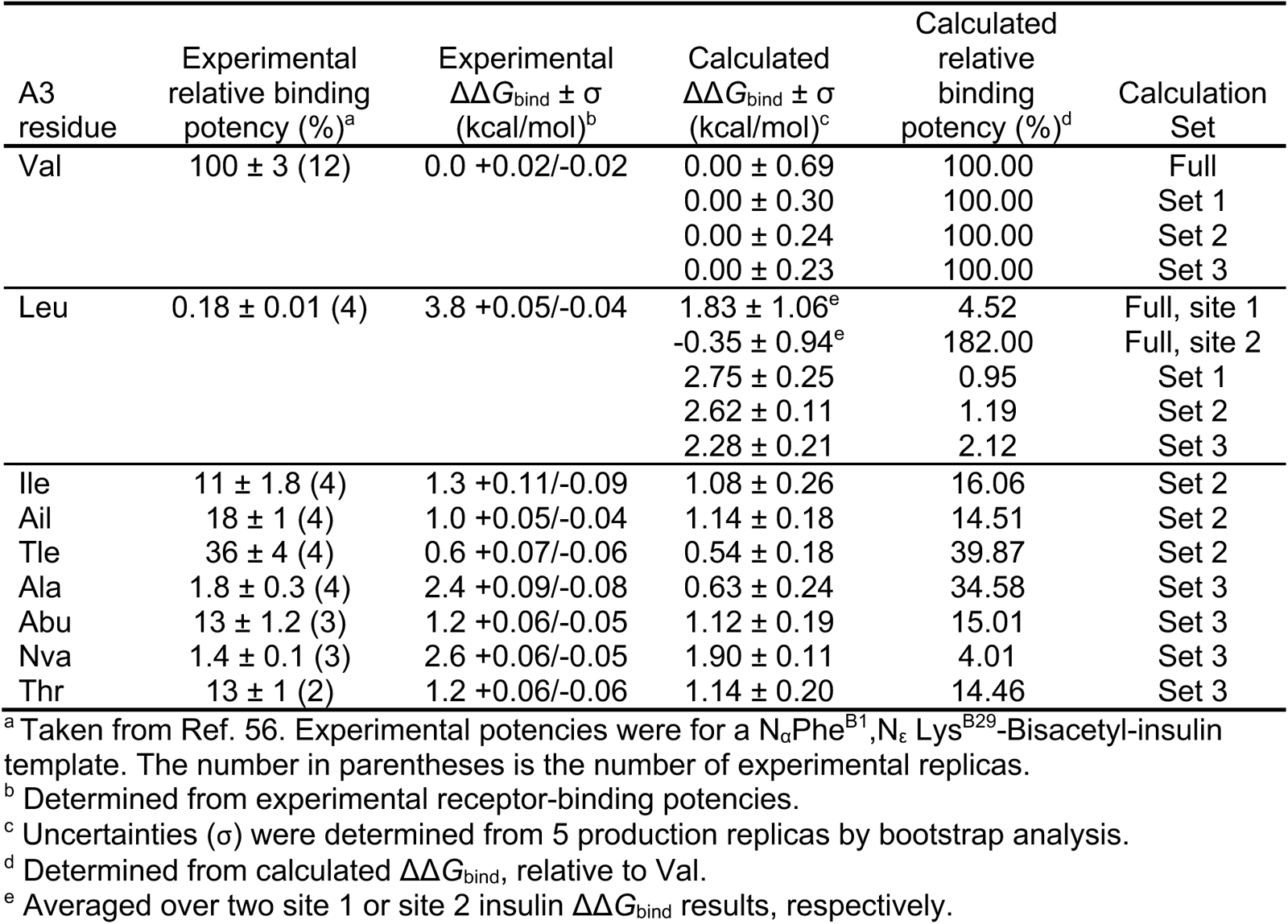
Experimental competitive binding affinities and calculated relative binding free energies (ΔΔ*G*_bind_) of eight insulin A3 variants.

To better focus simulation sampling on site 1 bound insulins, the full insulin-IR complex was spherically truncated around one IR site 1 with insulin bound (see Methods and Fig. S2). This reduces the size and complexity of the full IR system and is faster to sample with MD. The Set 1 Leu^A3^ mutation was then reevaluated with λD, which yielded a ΔΔ*G*_bind_ of 2.75 ± 0.25 kcal/mol, equivalent to a 100-fold loss of binding (Table 1, “Set 1”). Compared to the previous Full ΔΔ*G*_bind_, this result is nearer the experimental ΔΔ*G*_bind_ of 3.8 kcal/mol. The observed improvement in accuracy for Set 1 likely stems from the fact that the truncated system is smaller and has a lower computational cost per nanosecond, enabling longer simulations of 25 ns of MD sampling to be performed and relevant degrees of freedom of IR site 1 dynamics to be better modeled over longer timeframes. The improvement in sampling is also notable in the reduced bootstrapped errors in Set 1 compared to Full (Table 1). As a result of the improved agreement with experiment, the truncated system was then used for Sets 2 and 3 λD simulations to investigate additional A3 variants.

As shown in Fig. 3 and Table 1, calculated ΔΔ*G*_bind_ results for Set 2 and 3 perturbations showed high agreement with experiment. Five out of eight A3 mutants had computed ΔΔ*G*_bind_ results within 0.3 kcal/mol of experiment. Nva was within 0.7 kcal/mol of experiment, and Ala^A3^ was more outlier with an error of 1.8 kcal/mol. Computed errors for Leu^A3^ varied within a range of 1.0-1.5 kcal/mol of experiment, but Leu^A3^’s computed ΔΔ*G*_bind_ maintained statistical consistency between all three independent calculations, suggesting adequate and consistent sampling between all data sets. Compared to experiment, the mean unsigned error (MUE) across all data sets is 0.68 kcal/mol and the Pearson correlation is 0.88. This MUE is well below the typical target of 1.0 kcal/mol agreement with experiment used to evaluate state-of-the-art alchemical free energy calculations (66,72–74). This high degree of agreement between the λD calculated ΔΔ*G*_bind_ and experimental potencies confirms that a thermodynamic explanation exists for A3 variants’ loss of IR binding. Accordingly, a structural rationale of binding trends should be contained within the λD trajectories to help rationalize these computed ΔΔ*G*_bind_ results. Potential structural causes of insulin binding losses were identified by analyzing conformational orientations and insulin-IR interactions for each A3 side chain individually. Side chain dihedral angles were measured to identify how each residue was oriented within IR binding site 1. Each variant’s effect on local insulin-IR interactions was then probed by measuring select distances between insulin’s A chain N-terminus (A^Nter^) and the IR’s αCT peptide.

**Figure 3.**
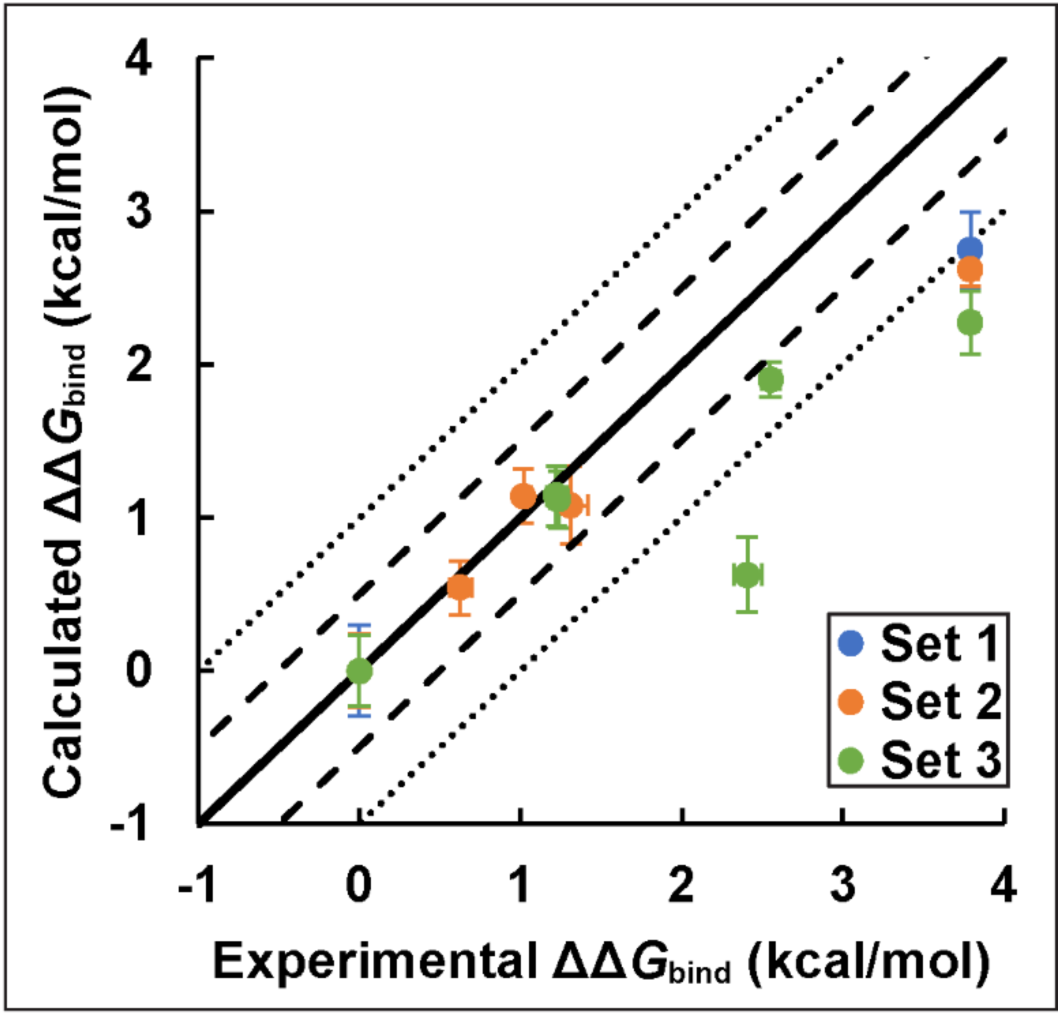
Correlation between experimental and λD computed relative binding free energies for eight insulin variants bound to the IR, relative to native insulin Val^A3^. Computed results were obtained over three calculation sets (Set 1 in blue, Set 2 in orange, and Set 3 in green). Ideal agreement between calculated and experimental ΔΔ*G*_bind_ results is represented by a solid black line. Dashed and dotted lines represent ΔΔ*G*_bind_ errors within ± 0.5 kcal/mol and ± 1.0 kcal/mol, respectively.

To investigate how the A3 side chains oriented within IR binding site 1, dihedral angles for all non-Ala side chains were calculated and clustered. Dihedral angle χ_1_ was measured between a residue’s N, C_α_, C_β_, and C_γ_ atoms. Additionally, for Leu, Ile, Ail, and Nva, dihedral angle χ_2_ was calculated between C_α_, C_β_, C_γ_, and C_δ_ atoms. As expected, frequency plots of one-dimensional χ_1_ (Fig. 4) or two-dimensional X_1_, X_2_ (Fig. 5) dihedrals showed clusters near 60°, 180°, or 300°, corresponding to stable gauge and trans conformations typically seen in small alkyl organic compounds. This natural conformational grouping allows λD frames to be sorted into bins of 0°≤X<120°, 120°≤X<240°, or 240°≤X<360° for analysis. In Fig. 4, bins are labeled 1-3 for Val, Tle, Abu, and Thr, and, in Fig. 5, bins are labeled 1-9 for Leu, Ile, Ail, and Nva. For most of the A3 variants, 70% or more of the variant frames are found in one or two dihedral bins, revealing clear preferences for specific side chain orientations within the insulin-IR complex. Fig. 4A shows that Val is conformationally locked into a χ_1_ = 180° orientation. This is notable because dihedral angles of alchemical functional groups are scaled by λ in λD, meaning barriers to rotation approach 0.00 kcal/mol as λ-scaling approaches zero. Previous studies have observed that this feature provides enhanced conformational sampling of alchemical perturbations in λD (67,75). But despite having the ability to rotate around χ_1_ when in an alchemically non-interacting state, Val^A3^ maintains the χ_1_ = 180° orientation in bin 2 in all calculation sets (Sets 1-3) whenever this residue is sampled by λD. Interestingly, Thr^A3^, which has similar C_β_ branching as Val^A3^, shows greater conformational flexibility and populates bins 2 and 3, with a slight preference towards bin 2. These trends reverse with Abu^A3^, which populates bin 3 (χ_1_ = 300°) more than bin 2. The symmetry of Tle^A3^ is nicely reproduced in Fig. 4D with three equally populated bins. Fig. 5A shows that Leu^A3^ populates bin 8 (χ_1_ = 300° and X_2_ = 180°) 58% of the time and bin 4 (χ_1_ = 180° and X_2_ = 60°) 37% of the time. Ail^A3^ similarly prefers two orientations, in bins 6 and 9, while Ile^A3^ and Nva^A3^ prefer a single orientation in either bin 6 or 9, respectively. Some additional flexibility is observed in Nva and Ail residues, however, these conformations were visited less than 10-15% of the time per bin respectively. Each bin’s frames were clustered and analyzed independently to identify representative structures and structurally depict dihedral angle trends observed in Figs. 4 and 5. In the analysis of Tle^A3^ and Ala^A3^, representative structures were identified from all production frames, since the symmetry of Tle^A3^ and the absence of a X_1_ in Ala^A3^ precludes frame clustering by dihedral angle. In the following discussions, the most and second-most populated dihedral bins are referred to as the primary and secondary dihedral clusters, respectively.

**Figure 4.**
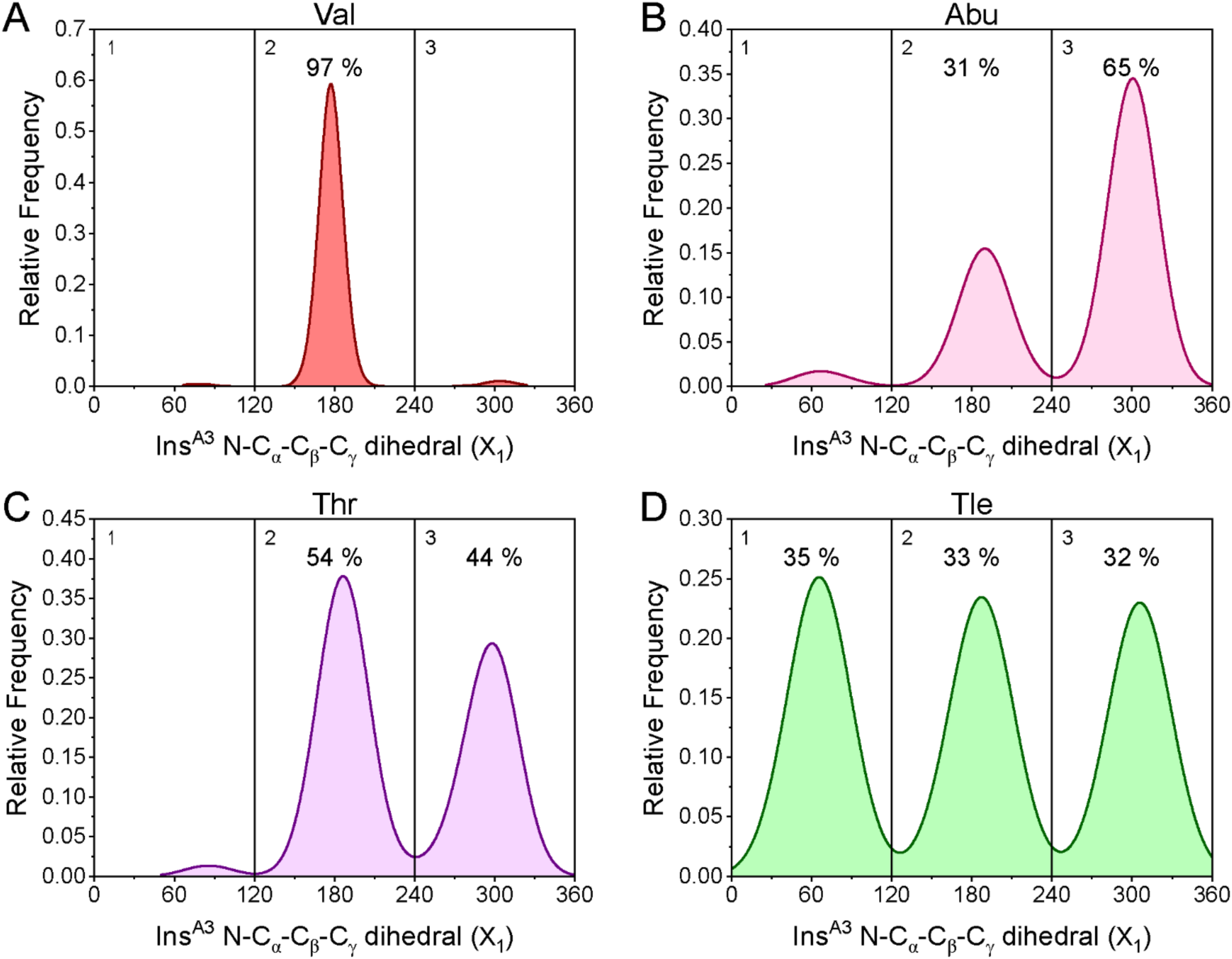
X_1_ Dihedral angle distribution plots of insulin A3 variants: Val (A), Abu (B), Thr (C), and Tle (D). Each plot is divided into bins of 120° and labeled 1-3. Bins populations are labeled if a bin contained 20% or more frames. Val^A3^ frames were combined from Sets 1-3.

**Figure 5.**
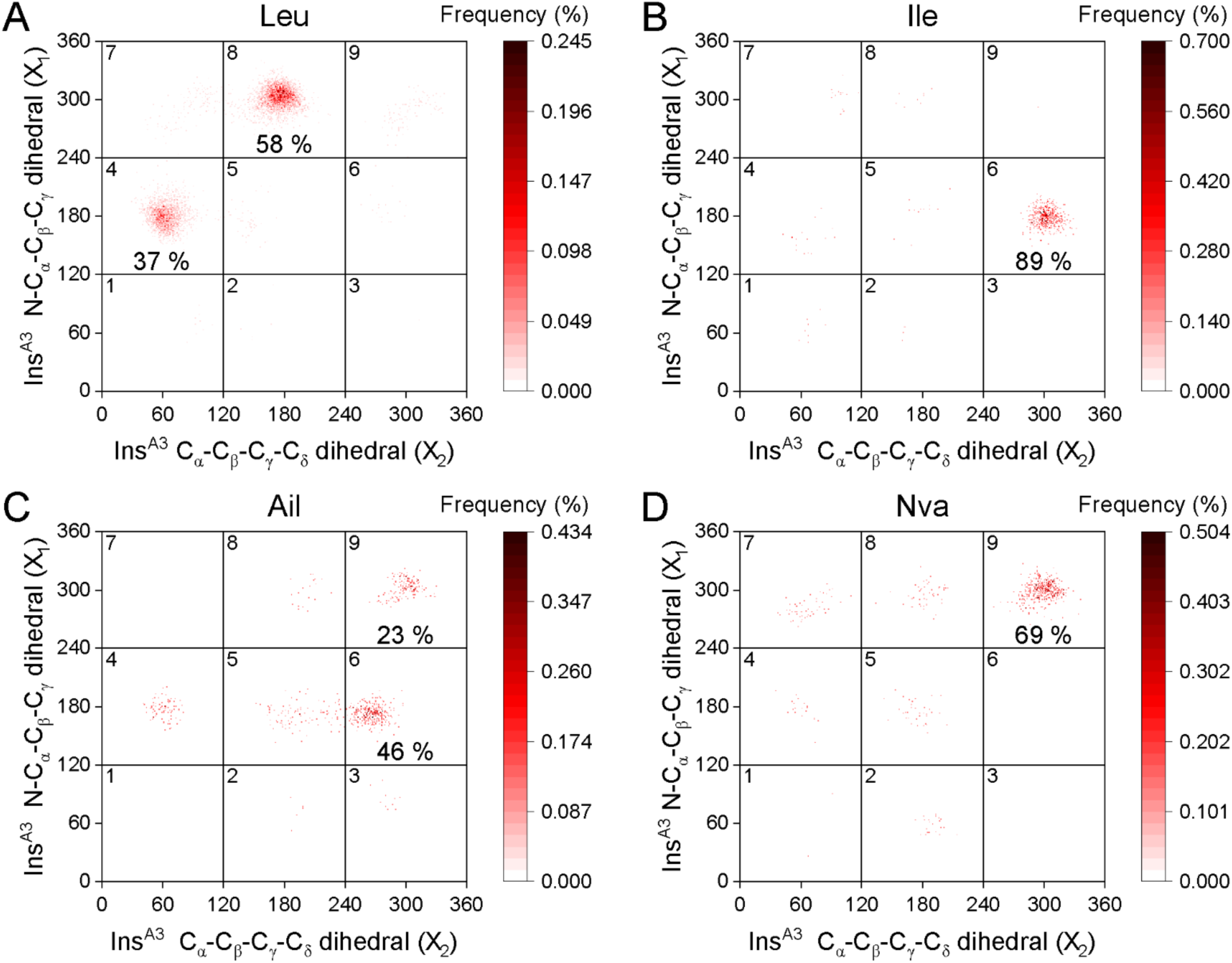
Two-dimensional X_1_, X_2_ dihedral angle distribution plots of insulin A3 variants: Leu (A), Ile (B), Ail (C), and Nva (D). Each plot is divided into bins of 120° per X angle and labeled 1-9. Bins populations are labeled if a bin contained 20% or more frames. Leu^A3^ frames were combined from Sets 1-3.

Representative frames were identified for primary and secondary dihedral clusters by computing probability densities for each side chain heavy atom in coordinate space and matching λD frames that best fit the collective probability densities (see Methods). As shown in Figs. 6 and S3, a majority of these probability density clouds clearly outline each side chain in one or two distinct positions. For instance, the heavy atom density for the primary Abu^A3^ cluster contains density fitting a single structure (Fig. 6G), but the secondary Abu^A3^ cluster shows density for two distinct structures (Fig. S3C). This was also observed for some Leu^A3^ secondary structures. Yet, for most residues and clusters analyzed, a single preferred conformation was observed. Across all three simulation sets, Val^A3^ clearly maintains a preferred orientation that keeps its C_γ2_ atom oriented to the right and its C_γ1_ atom oriented down and back (Fig. 6), as viewed from the IR-binding interface along insulin’s A^Nter^ (Fig. 7A). This orientation is similarly maintained in the primary dihedral clusters of all other β-branched A3 variants, such as Ile, Ail, Tle, and Thr (Fig. 6). In some secondary cluster orientations, such as Ail^A3^ and Thr^A3^, the side chain rotates so that both γ heavy atoms are pointed out towards the αCT peptide. Notably, just as Val^A3^’s C_γ1_ atom is partially solvent exposed, this secondary orientation allows Thr^A3^’s hydroxyl group to rotate out of the hydrophobic core and interact with solvent. Interestingly, both primary and secondary structures preferentially position one of its A3 side chain heavy atoms into the space where Val^A3^ keeps its C_γ2_ atom, which is in direct contact with the αCT peptide and occupies the cleft directly between Asp707, His710, and Asn711. As shown in Fig. 7, which highlights Val^A3^’s C_γ2_ atom density cloud aligned with all other side chain representative structures, all A3 residues fill this space in the insulin-IR complex. The one exception is Abu^A3^’s secondary representative structure, which has an alternative conformation and places its only C_γ_ atom in the position occupied by Val^A3^’s C_γ1_ atom (Fig S3C). Fig. S4 in the Supporting Information also shows aligned structures between each side chain and a representative Val^A3^ structure. High conformational similarities are observed for both small and large A3 mutations alike, and it is clear that each A3 variant adopts a Val-like conformation whenever possible. This tendency is most notable in Ala^A3^, Abu^A3^, and Nva^A3^ side chains which are less bulky and have more flexibility to reorient within a crowded space yet still predominantly positioned a heavy atom similar to Val^A3^. Importantly, for Leu^A3^ to maintain the placement of one of its C_δ_ atoms near Val^A3^’s C_γ2_ atom, severe movement in insulin’s A^Nter^ helix must occur, compared to Val^A3^’s A^Nter^ helix position. This A^Nter^ helical movement is largest for Leu^A3^ and likely a major contributing factor to its lowest insulin binding affinity among all eight A3 variants tested here. To further quantify conformational changes induced by the A3 mutations, nearby insulin-IR interaction distances were measured.

**Figure 6.**
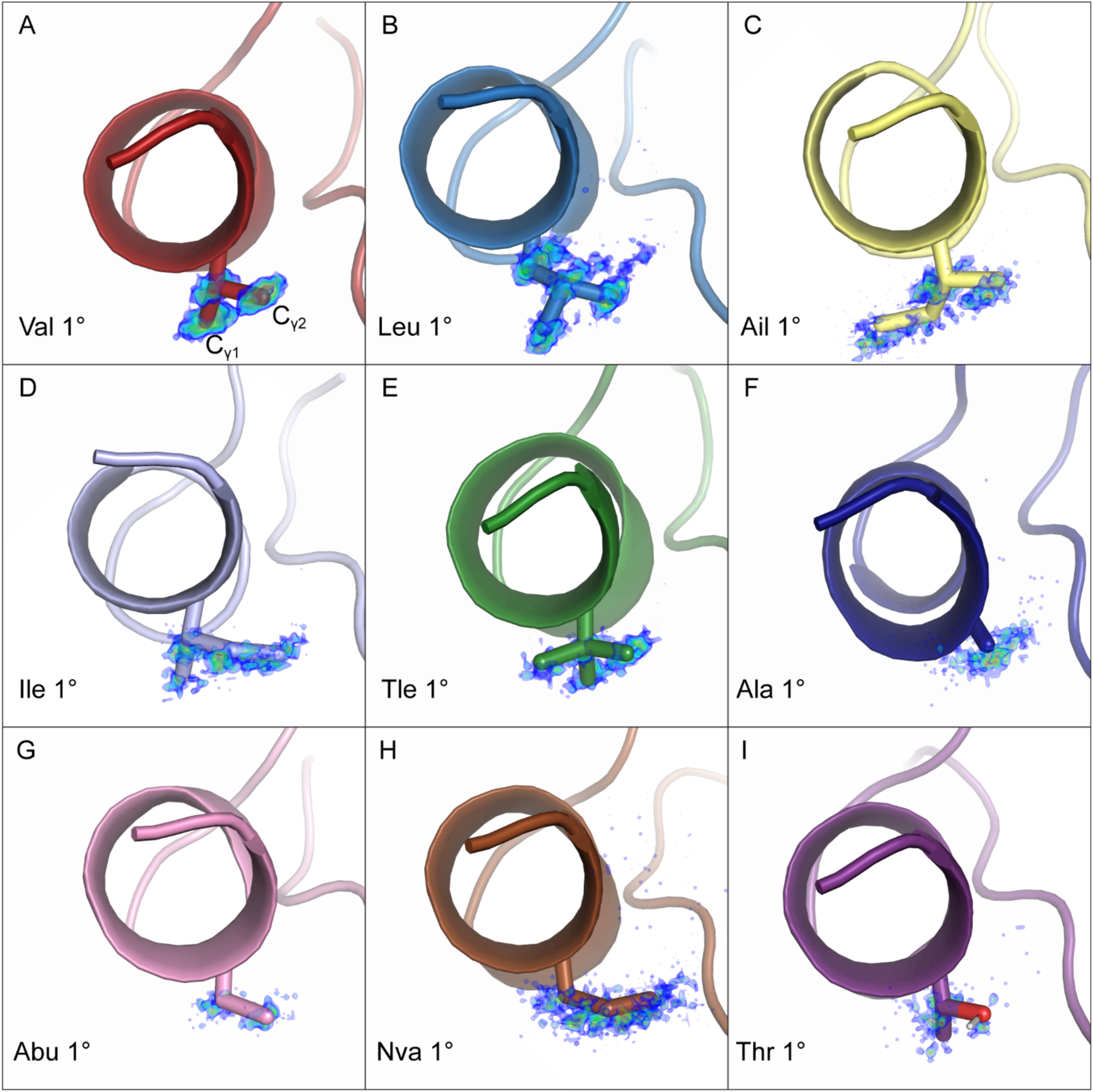
Representative frames for insulin A3 variant primary clusters: Val (A), Leu (B), Ail (C), Ile (D), Tle (E), Ala (F), Abu (G), Nva (H), Thr (I). A3 side chains are shown as stick representations, extending from insulin’s A-chain terminal helix. Side chain heavy atom probability densities are represented on a rainbow color scale from lower probability density (blue) to higher probability density (red).

**Figure 7.**
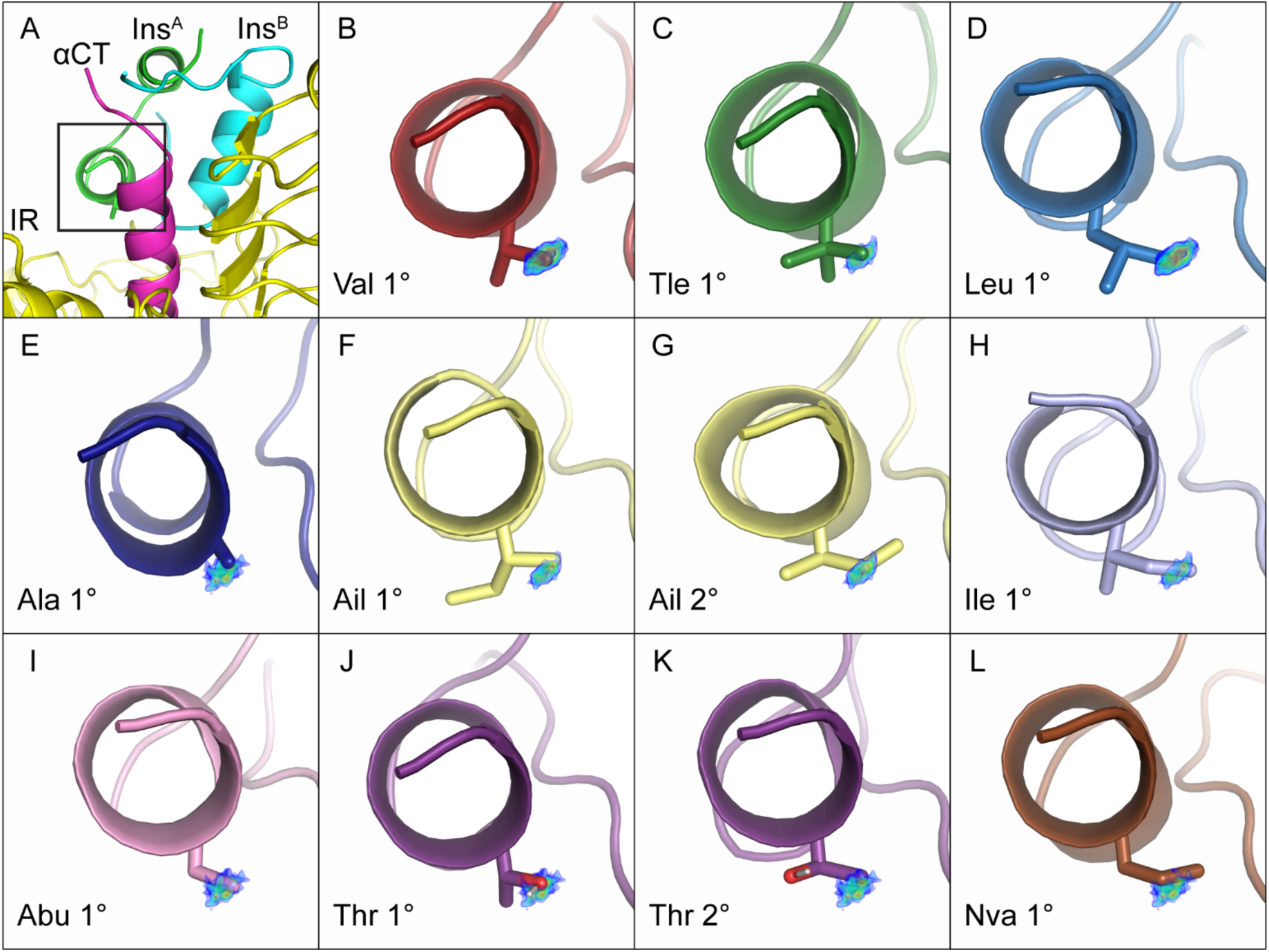
(A) Insulin bound at IR site 1. The black box shows the section and perspective of these images. The αCT peptide is hidden to allow full view of all insulin side chains. (B-L) Representative frames of insulin A3 variant primary (1°) and secondary (2°) clusters aligned with Val^A3^’s C_γ2_ probability density. Probability density is represented on a rainbow color scale from lower (blue) to higher (red) density.

Along the insulin-IR complex interface, the αCT peptide residue Asn711 is closest to insulin’s A3 residue and its carboxamide group is within hydrogen bonding distance to the A3 backbone amide NH or a neighboring A^Nter^ Glu^A4^ side chain carboxylate. To quantify potential changes to interactions between these residues upon A3 mutation, Asn711 X_1_ and Glu^A4^ X_2_ dihedral angles were measured along with Asn711 C_α_ – A3 N, Asn711 N_δ_ – Glu^A4^ O_ε_, and Asn711 O_δ_ – A3 N distances (labeled d_1_, d_2_, and d_3_ respectively, see Fig. 8A). Measurements were compiled from each primary and secondary dihedral cluster and plotted as violin plots to show the relative frequency of each dihedral or distance measurement (Figs. 8 and S5). In its preferred orientation, Val^A3^ maintains an average backbone N to Asn711 C_α_ distance of approximately 5.5 Å (Fig. 8B). This distance can be considered a relative measure of the insulin A^Nter^ proximity to the αCT peptide. As expected, this distance is shorter for smaller A3 variants (e.g. Ala^A3^) and longer for bigger side chains, including Ile^A3^, Ail^A3^ (2°), Nva^A3^, and Leu^A3^. The shorter distance for Ala^A3^ occurs because the insulin A^Nter^ helix moves forward and upward towards the αCT peptide to position Ala’s C_β_ where Val keeps its C_γ2_ atom (Fig. S4). Residues that adopt a Val-like conformation or are similarly sized, including the primary clusters of Abu^A3^, Ail^A3^, or Thr^A3^, maintain a median Asn711 C_α_ – A3 N distance consistent with Val^A3^. When there is a second C_γ_ oriented towards the αCT peptide (as in Tle^A3^ and the secondary clusters of Ail^A3^ and Thr^A3^) or when an A3 side chain has a C_δ_ positioned in insulin’s core (as in the Ail^A3^ secondary cluster, Ile^A3^, and Nva^A3^), the insulin A^Nter^-IR distance is modestly lengthened (Figs. 8B and S5B). Leu^A3^, on the other hand, has the most striking effect and lengthens this distance by almost 1.0 Å, in agreement with large structural distortions to insulin’s A^Nter^ chain observed in Figure S4.

**Figure 8.**
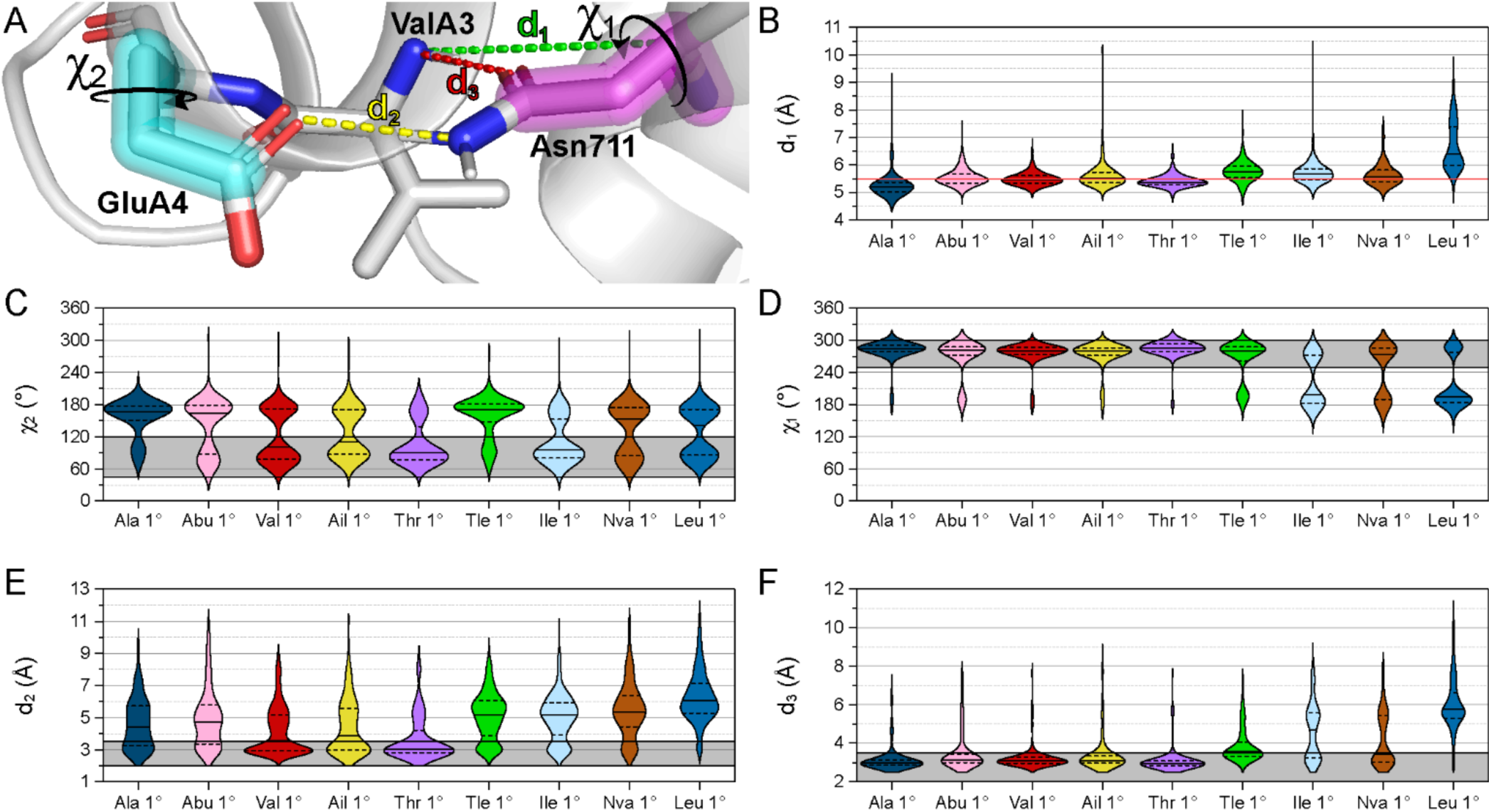
Insulin A^Nter^ and Asn711 distance and dihedral measurements for insulin A3 variant primary clusters. All violin plots are normalized to have the same area; solid lines represent median values and dashed lines show the interquartile range. (A) Structural depiction of distance and dihedral measurements in the insulin-IR complex. Insulin Glu^A4^’s X_2_ is highlighted in cyan. IR Asn711’s X_1_ is highlighted in pink. Distances are shown as colored dashed lines. (B) IR Asn711 C_α_ – Insulin A3 N atom distance (d_1_). The red line shows Val^A3^’s median value. (C) Insulin Glu^A4^ X_2_. The gray bar shows the dihedral window that facilitates hydrogen bonding between Asn711 and Glu^A4^ distance (d_2_). (D) IR Asn711 X_1_. The gray bar shows the dihedral window that facilitates hydrogen bonding to insulin A4 and A3 residues (d_2_ and d_3_, respectively). (E) IR Asn711 N_δ_ – Insulin Glu^A4^ O_ε_ distance (d_2_). The gray bar highlights distances below 3.4 Å. (F) IR Asn711 O_δ_ – Insulin A3 N distance (d_3_). The gray bar highlights distances below 3.4 Å.

Further analysis reveals insulin A^Nter^-IR hydrogen bonds are broken when insulin A3 variants clash with insulin Glu^A4^ or IR Asn711 side chains. Clustered frames for native Val^A3^ show that Glu^A4^ flips between two side chain orientations of χ_2_ ≈ 80° or 175° (Fig. 8C), while Asn711 rigidly maintains a consistent side chain conformation of χ_1_ ≈ 270° (Fig. 8D). Due to the rigidity of the Asn711 χ_1_ and the flexibility of Glu^A4^’s χ_2_, the Glu^A4^ and Asn711 side chains are only within hydrogen bonding distance of one another about half the time for Val^A3^ (Fig. 8E), while a hydrogen bond between Asn711 and the A3 backbone NH is nearly constant (Fig. 8F). For frames where the Asn711 N_δ_ and the Glu^A4^ O_ε_ atoms were within hydrogen bonding distance (between 2.0-3.4 Å, Fig. 8E gray bar), Glu^A4^’s χ_2_ was consistently between 45 and 120°, indicating a preferred conformation for Glu^A4^ to hydrogen bonding with Asn711. Gray bars in Fig 8 violin plots denote preferred Asn711 or Glu^A4^ conformations that encourage hydrogen bonding for Val^A3^ insulin, to make comparisons with other A3 variants easier. A3 side chains that are smaller than Val (Ala and Abu) or which orient two C_γ_ atoms towards the αCT peptide (Tle^A3^ and the secondary structures of Ail^A3^ or Thr^A3^) decrease the amount of time that Glu^A4^ has an appropriate dihedral angle to hydrogen bond with Asn711, resulting in lower likelihoods of forming Asn711 N_δ_ to Glu^A4^ O_ε_ hydrogen bonds (Fig. 8E). Additionally, the Tle^A3^ and secondary Ail^A3^ and Thr^A3^ structures also slightly perturb Asn711’s conformation (χ_1_ < 240°) and lengthen Asn711 O_δ_ – A3 N distances, reducing the likelihood of hydrogen bonding to between 50% and 30% of the time for these clusters. The largest change to Asn711 dihedral angle (Fig. 8D) comes from A3 side chains containing longer C_δ_-containing alkyl chains (Ile^A3^, Nva^A3^, and Leu^A3^). All three residues show large shifts in Asn711’s χ_1_ and corresponding changes in Asn711 O_δ_ – A3 N distances that weaken hydrogen bonding between insulin and the αCT peptide (Fig. 8F). This effect is largest for Leu^A3^, which directly clashes with Asn711 and breaks the Asn711 O_δ_ – A3 NH hydrogen bond more than 95% of the time. Notably, Asn711 N_δ_ – Glu^A4^ O_ε_ distances are also much longer for Leu^A3^ (Fig. 8E), indicating that almost no hydrogen bonding interactions exist for insulin *Wakayama* in the local pocket surrounding insulin’s A3 residue. Compared to trends observed for Val^A3^, a combination of steric clashes and the loss of 1-2 hydrogen bonds for Leu^A3^ would be expected to severely reduce binding affinity for insulin *Wakayama*.

## DISCUSSION

The results above describing binding orientations, representative structures, and relative changes in binding free energies for eight insulin A3 variants collectively illustrate how to maximize insulin’s A3 residue’s binding affinity to IR site 1. For it to have the most favorable binding affinity, insulin’s A3 residue should be branched off the C_β_ and orient one branched C_γ_ atom downwards and away from the αCT peptide while the other C_γ_ atom is positioned forward and to the right, matching the λD preferred conformation of Val^A3^. When the A3 side chain is modified such that it can no longer fulfill both of these requirements, a preference emerges for A3 side chains to fill the space occupied by Val’s C_γ2_ atom (Fig. 7). As a result, disruptive conformational changes may occur which introduce steric clashes and disrupt favorable hydrogen bonding interactions, weaking insulin’s IR binding affinity.

To further understand these effects, patterns in computed ΔΔ*G*_bind_ between A3 mutants can be decomposed into a per-carbon-unit (CH_n_) basis. The impact on insulin’s binding affinity for IR site 1 due to the loss of Val^A3^’s C_γ1_ atom can be estimated by comparing Abu^A3^ versus Val^A3^ results. Primary structures for both residues (Fig. 7I and B, respectively) and their effects on local dihedral angles or distances (Fig. 8) show that these two residues occupy very similar conformations and that Abu does not generally disrupt insulin’s local interactions with the IR, yet insulin Val^A3^’s calculated binding affinity is 6.7x larger than Abu^A3^’s (Table 1). Similarly, when the unbranched Nva^A3^ is compared to the β-branched Ile^A3^, primary structures (Fig. 7L and H, respectively) and local dihedral angle and distance trends are very similar, yet insulin Ile^A3^’s calculated binding affinity is 4.0x larger than insulin Nva^A3^’s. Combined, this suggests that losing a C_γ_ atom in the A3 side chain results in approximately a 4-to-7-fold loss in insulin-IR binding affinity, which predominantly correlates to the loss of Val^A3^’s C_γ1_ atom. Notably, the striking similarity between Ile^A3^ and Nva^A3^’s primary structures and local dihedrals/distance trends are not due to the two variants influencing each other during the λD simulations because these two A3 variants were sampled in separate simulations.

An analogous comparison of the same four insulin A3 variants can also provide ΔΔ*G*_bind_ insights into the impact of adding a C_δ_ atom to insulin’s A3 residue. Comparing Ile^A3^ and Val^A3^ (both β-branched but Ile^A3^ contains an additional C_δ_ atom) or Nva^A3^ and Abu^A3^ (both unbranched but Nva^A3^ contains an additional C_δ_ atom) reveals that while both sets of side chains are similarly oriented in IR site 1, the addition of an extra C_δ_ atom has a more pronounced effect on local environmental dihedral angles and insulin-IR distances than the loss of the C_γ1_ atom. In both Ile^A3^ and Nva^A3^, the added C_δ_ atoms clash with Asn711 and lengthen the distance between the insulin A^Nter^ and the IR, diminishing the frequency of Asn711-A^Nter^ hydrogen bonds to both insulin A3 and A4 residues. As Val^A3^ has 6.2x the binding affinity of Ile^A3^ and Abu^A3^ has 3.7x the binding affinity of Nva^A3^, this suggests a similar approximate 4-to-6-fold loss in insulin-IR binding affinity due to the addition of an A3 C_δ_ atom. Finally, for comparing Tle^A3^ and Val^A3^, the addition of an extra C_γ_ atom only contributes to ca. 2.5-fold loss in insulin binding. For Tle^A3^, while the extra C_γ_ clashes with Glu^A4^ and reduces Tle^A3^ binding compared to Val^A3^, it has a much smaller effect on Asn711’s interactions and conformations.

The loss of affinity of Ail^A3^ and Thr^A3^ is less straightforward to decompose energetically due to their inconsistent population of two dihedral clusters and orientations. Both Ail^A3^’s and Thr^A3^’s primary structures have remarkably similar orientation and local dihedral angles and distances as Val^A3^ (Figs. 7, 8, and S4). These orientations are the computed preferred states for both Thr^A3^ and Ail^A3^, despite Thr^A3^’s hydroxyl sticking into a hydrophobic pocket (albeit still within hydrogen bonding distance of Asn711 which it is regularly observed interacting with) and Ail’s longer chain sticking into solvent near the A3 binding pocket. In their secondary orientations, however, Ail^A3^ and Thr^A3^ both lose the heavy atom filling Val^A3^’s C_γ1_ pocket (analogous to Abu^A3^ and Nva^A3^ discussed above) and have a second C_γ_ pointing towards the αCT peptide and clashing with Glu^A4^ (similar to Tle^A3^). Estimates above suggest both secondary configurations would have much less favorable binding affinities than Val^A3^, on an estimated order of ca. 10-17.5-fold. However, these residues have only 6.9-fold less computed affinities than Val^A3^ due to the higher weight and favorability of their primary structure on binding (Figs 4 and 5). As with Ile^A3^ and Nva^A3^, similarities in structure and local dihedral and distance trends for both Ail^A3^ and Thr^A3^ residues were captured across two different λD simulations.

Finally, as mentioned above, Leu^A3^ induces the most striking changes to insulin binding. It breaks both Asn711 hydrogen bonds with the A^Nter^ and forces the A^Nter^ away from the αCT peptide. This occurs because of its C_γ_ branching. If the primary structures of Leu^A3^, Ile^A3^, and Nva^A3^ are compared, all three residues point their C_γ_ atom towards the αCT peptide and the C_δ_ atom of their longest chain back towards insulin’s core (Figs. 6 and S3). Leu^A3^, however, has an additional C_δ_ atom that can only be positioned towards the αCT peptide regardless of its side chain orientation. Both primary and secondary structures of Leu^A3^ reflect this (Figs. 6 and S3), and energetically, Leu^A3^’s computed potency is 47.4-105.3-fold less than Val^A3^ (Table 1, Sets 1-3). In an approximate estimate, this correlates to the minimum fold losses described above of concurrently losing one C_γ_ atom and gaining two C_δ_ atoms (4 × 4 × 4 = 64-fold loss). Thus, Leu^A3^’s unique structural composition as a hydrophobic side chain, specifically its γ-branching and extra C_δ_ atom, introduces steric clashes with the IR’s αCT peptide, substantially worsening insulin IR binding affinity.

In conclusion, insulin *Wakayama* and several other insulin A3 variants were characterized thermodynamically and structurally with λD alchemical free energy calculations. λD-based simulations are uniquely posed to compute binding affinity losses caused by chemical changes to a protein complex, such as a missense mutation to one or both protein partners and can model multiple mutations simultaneously within a single MD simulation. In addition to computing a quantitative ΔΔ*G*_bind_, λD trajectory frames can be analyzed to provide a structural rationale of computed free energy trends among investigated mutants. In this work, eight A3 variants, including the *Wakayama* Leu^A3^, were investigated and compared to native Val^A3^ insulin. Relative binding free energies were calculated among three sets of A3 mutations spanning both large and small perturbations. The sampling dynamics of each A3 side chain were characterized by 1-2 dihedral angles, which facilitated clustering of λD frames and the identification of preferred representative structures for each variant. The impact of each variant on insulin-IR interactions were then analyzed on a per cluster basis. The investigations into the structural rationale behind these observed binding trends revealed that local changes around the A3 mutation site are responsible for weaker binding of all the A3 variants. For example, our observations clearly delineate why Val alone is tolerated at insulin’s A3 position when insulin binds the IR. Valine has β-branching of two hydrophobic moieties that fit perfectly within the IR binding site 1, adjacent to Asn711. Val^A3^’s short size, compared to Leu^A3^, maximizes both favorable hydrophobic interactions within the pocket and encourages hydrogen bonding interactions between insulin A3 or A4 residues and Asn711 in the IR αCT peptide. Both larger and smaller residues disrupt some element of this fit, yielding weakened variant binding affinities. Leu^A3^ shows the largest structural changes with corresponding largest reduction in binding affinity. With side chain branching at its C_γ_ atom, severe steric clashes with IR’s αCT peptide and Asn711 are observed with Leu^A3^’s second C_δ_ atom. This results in insulin’s A^Nter^ being pushed away from the IR and breaks of almost all hydrogen bonding interactions between insulin and Asn711. Thus, this work confirms Nakagawa and Tager’s longstanding hypothesis that C_γ_ branching in leucine is central to insulin *Wakayama*’s 140-500-fold worse binding affinity. This work provides new atomistic insights and structural rationale clarifying this decades-old mystery. Furthermore, we note that structural patterns of A3 mutations were produced with high reproducibility across different λD simulations. Therefore, this work demonstrates the power to investigate critical PPIs with λD, both within the insulin-IR complex and more broadly in other protein-protein complexes.

## METHODS

The fully populated cryo-EM insulin–IR complex (PDB: 6PXW) was used as a structural starting point for λD modeling of the Full system (26). Hydrogen atoms were added to the structure using MolProbity, which also checks for histidine, asparagine, and glutamine side chain flips to optimize potential hydrogen bonding interactions (76,77). Missing internal loops in the 6PXW IR structure were repaired by splicing in coordinates from alternative IR structures (PDBs: 6SOF or 6HN5), when corresponding loops were resolved in those structures (17,27). Minimization was then performed in CHARMM using the steepest descent algorithm to reduce potential steric clashes with inserted residues. Histidine protonation states (His-δ or His-ε) were determined by manual inspection of the structure to identify potential hydrogen bonding partners and compared to MolProbity predictions determined with the Reduce program (78). PROPKA was used to identify potential protonation state changes to Asp, Glu, Lys, Arg, and His residues (79). For λD calculations Sets 1–3, spherical truncation of the repaired IR around a site 1 bound insulin (chain C in 6PXW) was performed by removing all protein residues outside a 27 Å radius centered on insulin from the IR structure. In λD simulations of the truncated system (described below), all protein backbone atoms outside a 17 Å radius from insulin were harmonically restrained with a force constant of 10 kcalmol^-1^Å^-2^ to maintain the IR’s local fold and shape during MD sampling. All other atoms in the chemical system remained fully flexible. To calculate relative free energies of binding with λD, alchemical perturbations must be performed in bound and unbound states (Fig. 2). Accordingly, a model of an insulin monomer was created by simply extracting coordinates for a site 1-bound insulin (chain C in 6PXW) into a separate pdb file.

Both full and truncated insulin-IR systems and the insulin monomer were solvated and neutralized in a 0.1 M NaCl solution using the CHARMM-GUI solution builder tool (80). A cubic box of TIP3P water was generated by ensuring that all box edges were at least 10 Å from a protein molecule on every side, and ions were added to ensure all systems had a neutral net charge with the desired buffer concentration (81). The CHARMM36 protein force field was used to represent all protein energetics (82). Prior to running λ-dynamics calculations, all protein molecules were harmonically constrained, and the solvent was minimized for 200 steps with a steepest descent algorithm. Constraints on protein atoms were then removed, and the entire system was minimized for another 250 steps. In the case of the truncated insulin-IR system, backbone harmonic restraints were retained for minimization and subsequent molecular dynamics.

λD was performed using the CHARMM molecular software package with the domdec module to facilitate use of graphic processing units (GPUs) (83–85). A cutoff value of λ ≥ 0.99 was used to classify perturbation end states. Simulations were run in the isothermal-isobaric ensemble with a temperature range of 25-30 °C to match clinical and experimental procedures and a pressure of 1.0 atm. MD simulations were run with a time step of 2 fs. Long-range interaction energies were modelled using particle mesh Ewald with force switching and nonbonded cutoffs between 9 and 10 Å (86–89). Hydrogens were restrained using the SHAKE algorithm (90).

λD simulations were performed as described previously for analyzing protein side chain perturbations (68,70,71). Briefly summarized, the Adaptive Landscape Flattening (ALF) algorithm was used to identify appropriate biasing potentials for each λ state in our λD simulations, which biases flatten the free energy landscapes in λ-space and enable the dynamic sampling of many different perturbation end states (65,91). Optimal biases were obtained from ALF after an aggregate of 20-45 ns of sampling. Five replica λD productions simulations were run for each bound and unbound system. All unbound insulin and the Full insulin-IR complex production simulations were run for 15 ns. Set 1 and 2 production simulations ran for 25 ns, and Set 3 production simulations were extended to 50 ns to obtain improved sampling. The first fifth of each production simulation was excluded as equilibration prior to free energy determination. Correspondingly, in the discussion below, the term “production frames” refers to frames sampled within the last 80% of a λD trajectory in any of the five replicas, i.e., after equilibration was removed. Final relative free energy differences (Δ*G*_P1_ or Δ*G*_P1P2_) were calculated using WHAM (92), and the final ΔΔ*G*_bind_ were obtained by taking the difference between bound and unbound results, as shown in Fig. 2. Statistical errors were estimated with bootstrapping.

Since the identity of the A3 residue changes dynamically throughout a λD simulation, the λD trajectory frames must be sorted by end state before structural analyses can be performed. An insulin frame was classified as belonging to one specific insulin variant if the λ value of an alchemical side chain was above a λ = 0.99 threshold, the same threshold used to compute Δ*G* differences. If none of the A3 variants have a λ value greater than 0.99 in a given frame, then the system was classified as a nonphysical intermediate state and the frame was discarded. After a frame was identified as a physically relevant insulin end state, the A3 side chain X_1_ and X_2_ dihedral angles were calculated with CHARMM. Where possible, each insulin variant’s production frames were then sorted into primary (1°) or secondary (2°) clusters based on their A3 side chain X_1_ and X_2_ dihedral angles (0°≤X<120°, 120°≤X<240°, or 240°≤X<360°). As the side chains of Ala and Tle (due to its symmetry) could not be sorted by A3 X_1_ and X_2_ dihedral angles, their primary clusters contain all the production frames for that variant.

Primary and secondary clusters were examined independently to determine representative structures for each cluster. The spatial distributions of each side chain’s alchemical heavy atoms were determined by calculating the probability of finding each heavy atom in a Cartesian coordinate grid. Similar to forming a two-dimensional histogram but in three-dimensional space, atomic positions were counted within cubic bins of 0.2 Å width in all directions and normalized by the number of frames for each cluster. The resultant probability density map could then be loaded into Pymol for visualization (93). To select a representative structure with the highest agreement matching the spatial distribution plot, the spatial distribution bins with at least 85% of the population of the maximum bin were collected and the weighted average was calculated. The frame that had the lowest deviation from the weighted average position for all side chain heavy atoms was selected as the representative frame for that density. When the variant dihedral clusters contained density for more than one structure (for example see the side chain densities and corresponding structures of Abu^A3^ 2° in Fig. S3C), a seed frame from the appropriate variant dihedral cluster that matched each density was first identified by visual inspection. These manually selected structures were then used as a guide to sort spatial distribution bins into belonging to one substructure or another (similar to sorting bins in a bi-modal histogram as belonging to one peak or another). After sorting, weighted average positions and corresponding representative structures were determined as described above. All structures and probability densities were visualized and imaged with PyMOL (93).

Finally, for every frame belonging to a primary and secondary structure, selected dihedral angle and distance measurements were determined. The IR Asn711 χ_1_ (N-C_α_-C_β_-C_γ_) and insulin Glu^A4^ X_2_ (C_α_-C_β_-C_γ_-C_δ_) dihedral angles and three distances of interest (the IR Asn711 C _α_-insulin^A3^ N distance, the insulin^A3^ N-IR Asn711 O_δ_ distance, and the distance between the IR Asn711 N_δ_ and the closer of the two insulin Glu^A4^ Oε atoms) were calculated using an in-house python script. We note that these measurements are also readily available with use of the CHARMM software suite. The distributions of these measurements within each primary and secondary cluster were then plotted as violin plots with OriginPro (94).

## ACKNOWLEDGMENTS

The authors gratefully acknowledge the National Institutes of Health (NIH), through grant R35GM146888, for financial support. The authors acknowledge the Indiana University Pervasive Technology Institute for providing supercomputing and storage resources that have contributed to the research results reported within this paper. We thank Dr. Michael A. Weiss for bringing this problem to our attention, for helpful discussions about this work, and review of a manuscript preprint.

## AUTHOR CONTRIBUTIONS

MPB: Conceptual design, data acquisition and analysis. JZV: Conceptual design, supervision, project administration, funding. All authors contributed to writing.

