## Supplemental Figures for "A λ-dynamics investigation of insulin *Wakayama* and other A3 variant binding affinities to the insulin receptor"

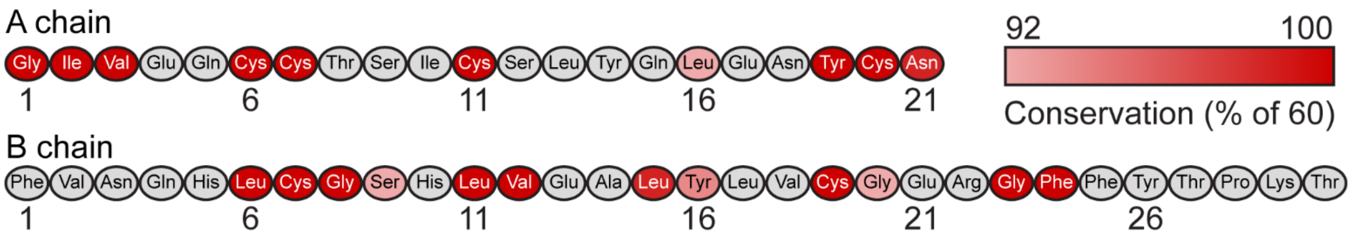


**Figure S1.** Human insulin sequence with color coded conserved residues. Residues that are conserved in at least 55 out of 60 species are colored with respect to their percent conservation. Conservation data is from reference: J. M. Conlon, Evolution of the insulin molecule: insights into structure-activity and phylogenetic relationships. *Peptides* **22**, 1183-1193 (2001).

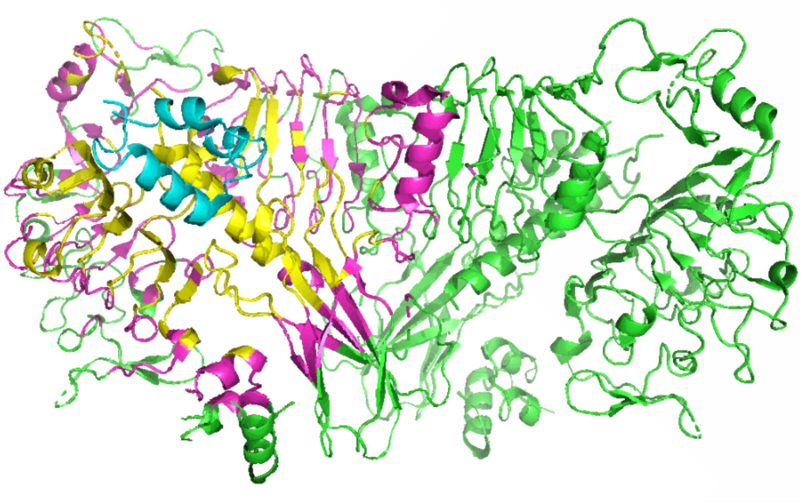

**Figure S2.** Truncated model of the insulin – insulin receptor complex. The ectodomain of the IR (PDBID: 6PXW; in green) was spherically truncated within 27 Å of the site 1 bound insulin (in cyan). IR residues that were retained are highlighted in magenta and yellow. Yellow residues were within a 17 Å radius of insulin and were fully flexible in the molecular dynamics simulation. Backbone atoms of magenta-colored residues were harmonically constrained to maintain the shape of the IR and its binding site.

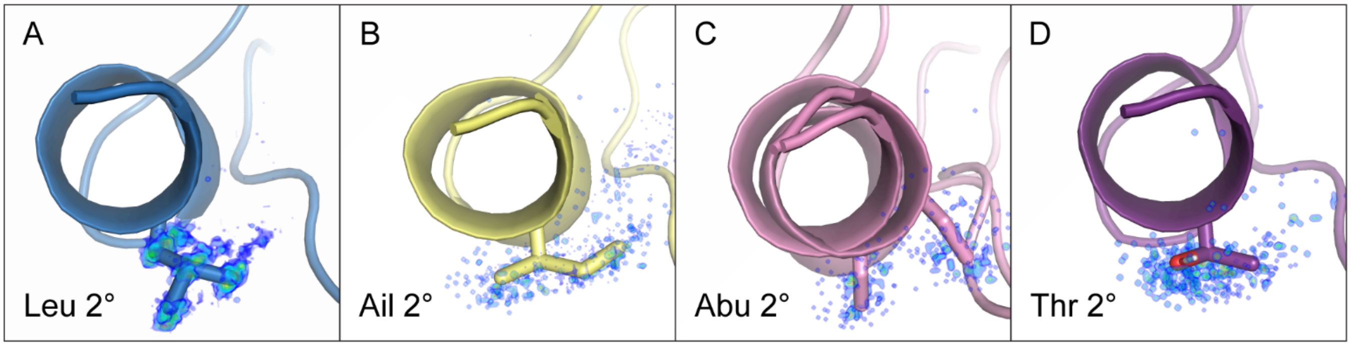


**Figure S3**. Representative frames for insulin A3 variant secondary clusters: Leu (A), Ail (B), Abu (C), and Thr (D). A3 side chains are shown as stick representations, extending from insulin's A-chain terminal helix. Side chain heavy atom probability densities are represented on a rainbow color scale from lower probability density (blue) to higher probability density (red).

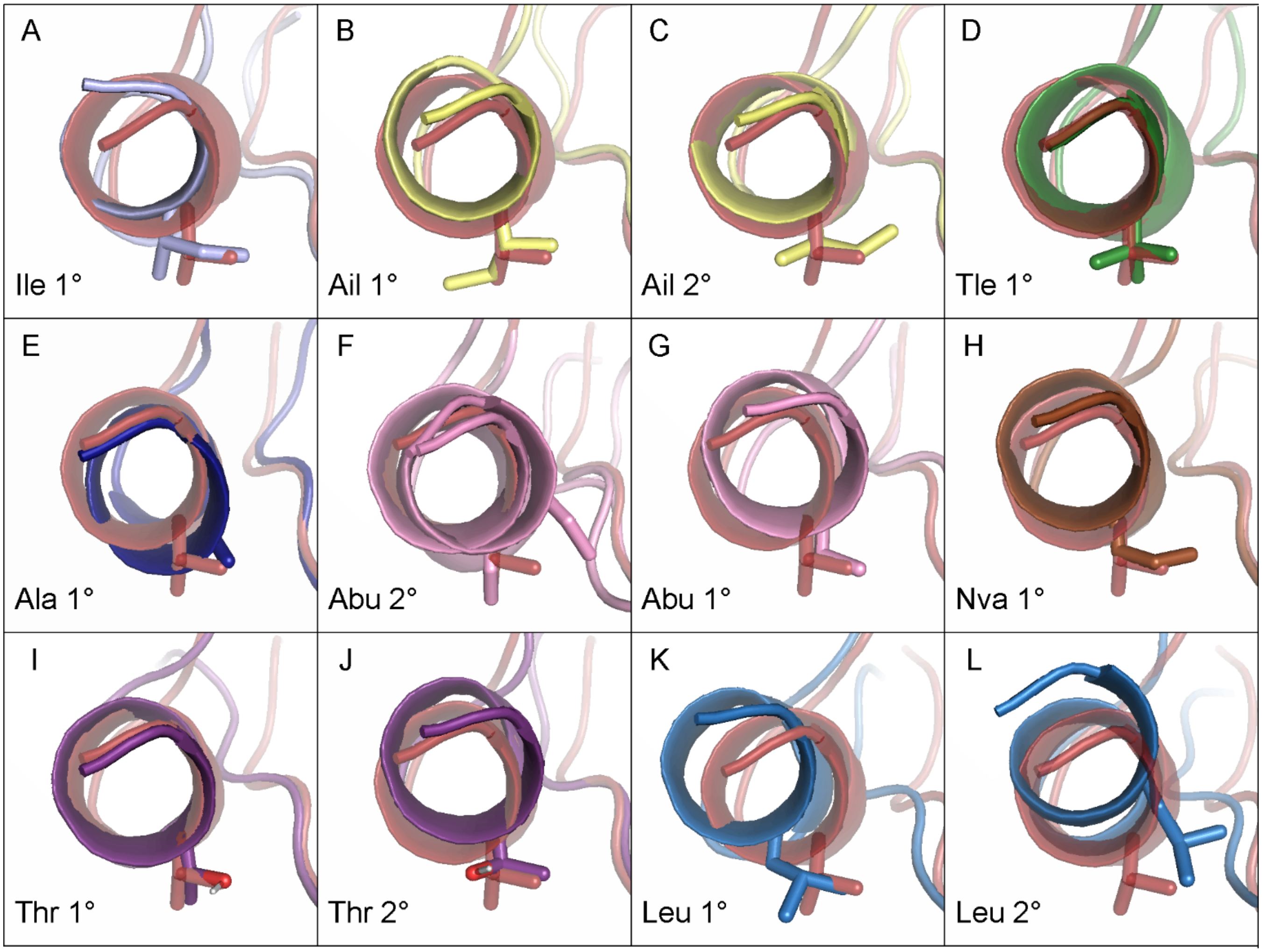


**Figure S4.** (A-L) Representative frames of insulin A3 variant primary (1°) and secondary (2°) clusters aligned with Val$^{A3}$'s primary representative frame in red (see Fig. 6A in the main text).

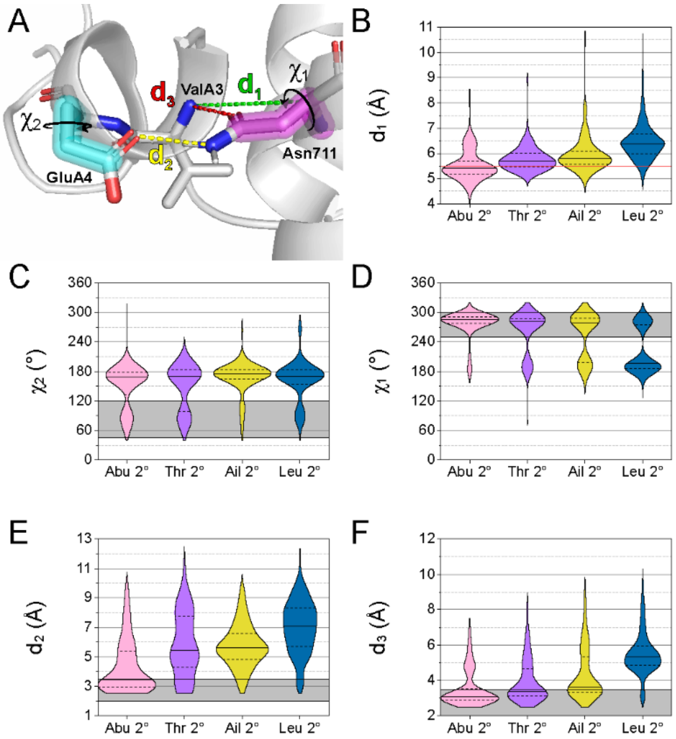


**Figure S5.** Insulin $A^{Nter}$ and Asn711 distance and dihedral measurements for insulin A3 variant secondary clusters. All violin plots are normalized to have the same area; solid lines represent median values and dashed lines show the interquartile range. (A) Structural depiction of distance and dihedral measurements in the insulin-IR complex. Insulin $Glu^{A4}$'s $\chi_2$ is highlighted in cyan. IR Asn711's $\chi_1$ is highlighted in pink. Distances are shown as colored dashed lines. (B) IR Asn711 $C_\alpha$ – Insulin A3 N atom distance ($d_1$). The red line shows $Val^{A3}$'s median value (from Fig. 8 in the main text). (C) Insulin $Glu^{A4}$ $\chi_2$. The gray bar shows the dihedral window that facilitates hydrogen bonding between Asn711 and $Glu^{A4}$ (distance $d_2$). (D) IR Asn711 $\chi_1$. The gray bar shows the dihedral window that facilitates hydrogen bonding to insulin A4 and A3 residues ($d_2$ and $d_3$, respectively). (E) IR Asn711 $N_\delta$ – Insulin $Glu^{A4}$ $O_\varepsilon$ distance ($d_2$). The gray bar highlights distances below 3.4 Å. (F) IR Asn711 $O_\delta$ – Insulin A3 N distance ($d_3$). The gray bar highlights distances below 3.4 Å.
